# Heterogeneous Effects of Src Inhibition on Determinants of Metastasis in Preclinical Models of Human Bladder Cancer

**DOI:** 10.1101/2024.11.09.622804

**Authors:** Kai Aragaki, Bryan Wehrenberg, Yujiro Hayashi, David J. McConkey

**Author notes:** These authors contributed equally (Kai Aragaki, Bryan Wehrenberg).

## Abstract

**Background:** Past work in preclinical models of solid tumors have implicated SRC in invasion and metastasis, but also demonstrated it inhibited bladder cancer metastasis.

**Objective:** Determine if the role of SRC in metastasis is dependent on bladder cancer molecular subtype membership.

**Methods:** We analyzed large public datasets, performed *in vitro* invasion and migration assays using small-molecule and doxycycline inducible SRC knock-down constructs, and *in vivo* experimental metastasis assays.

**Results:** Looking at large public datasets, we found SRC is upregulated in luminal papillary muscle invasive bladder cancer. Using the consensus classifier on RNA expression from 30 cell lines, we demonstrated that chemical SRC antagonists inhibited migration in luminal papillary cells but had little effect in basal/squamous lines. Conditional SRC knockdown inhibited migration in luminal papillary RT112 cells, whereas it increased migration and reduced proliferation in luminal papillary UM-UC6 cells. Regardless, these effects did not affect levels or sites of experimental metastasis *in vivo*.

**Conclusions:** The results support the conclusion that SRC’s biological effects in bladder cancer are not primarily involved in promoting invasion and metastasis. Further work is required to define SRC’s roles in luminal papillary bladder cancers.

## INTRODUCTION

The SRC protooncogene was first identified as the causative factor that mediated the transforming effects of an avian leukemia virus^1^. Its human counterpart encodes a 60kD non-receptor tyrosine kinase^2^ that has been implicated in cell cycle progression and integrin-mediated adhesion signaling. Early studies in hematopoietic cells and solid tumor cell lines implicated the integrin-dependent effects of SRC in directional motility, invasion, and metastasis^3^. However, studies in preclinical models of bladder cancer concluded that SRC inhibited invasion and metastasis via mechanisms that involved direct phosphorylation of RhoGDI2 and downstream inhibition of caveolin-1^4^,^5^.

Bladder cancers are highly heterogeneous in their invasive and metastatic potentials. The majority of bladder cancers are superficial low-grade papillary lesions (non-muscle-invasive bladder cancer, NMIBC) that are prone to recurrence but rarely progress to become life-threatening and metastatic. However, approximately 20-25% of bladder cancers are muscle-invasive at diagnosis, and about half of patients with muscle-invasive bladder cancer (MIBC) ultimately die of metastatic disease^6^. MIBC can also be grouped into basal and luminal molecular subtypes that are similar to those described in breast cancer^7–9^. Luminal bladder cancers are enriched with activating FGFR3 mutations and fusions^10^,^11^, higher expression of differentiation markers, and better survival outcomes^7^, whereas basal bladder cancers exhibit features of epithelial-to-mesenchymal transition (EMT) and are associated with invasive and metastatic disease at clinical presentation and shorter disease-specific survival^6^,^7^. We hypothesized that SRC plays different roles in luminal versus basal bladder cancers. We tested this hypothesis using public human bladder cancer bulk RNA expression datasets, human bladder cancer cell lines, and *in vivo* metastasis models.

## RESULTS

### SRC expression across subtypes

We first used public bulk mRNA expression datasets to explore if SRC expression correlated with stage and molecular subtype membership. Consistent with previous findings^12^, SRC expression was significantly higher in NMIBC relative to MIBC (Figure 1, A). SRC levels were relatively high in the least aggressive UROMOL subtype (Class 1), but also in the subtype associated with the highest rate of progression (Class 2a) (Figure 1, B)^13^. However, SRC expression strongly correlated with progression score in Class 1 but not Class 2a tumors, (Figure 1, C), implying SRC expression may play an indirect role in muscle invasion by potentiating progression.

**Figure 1:**
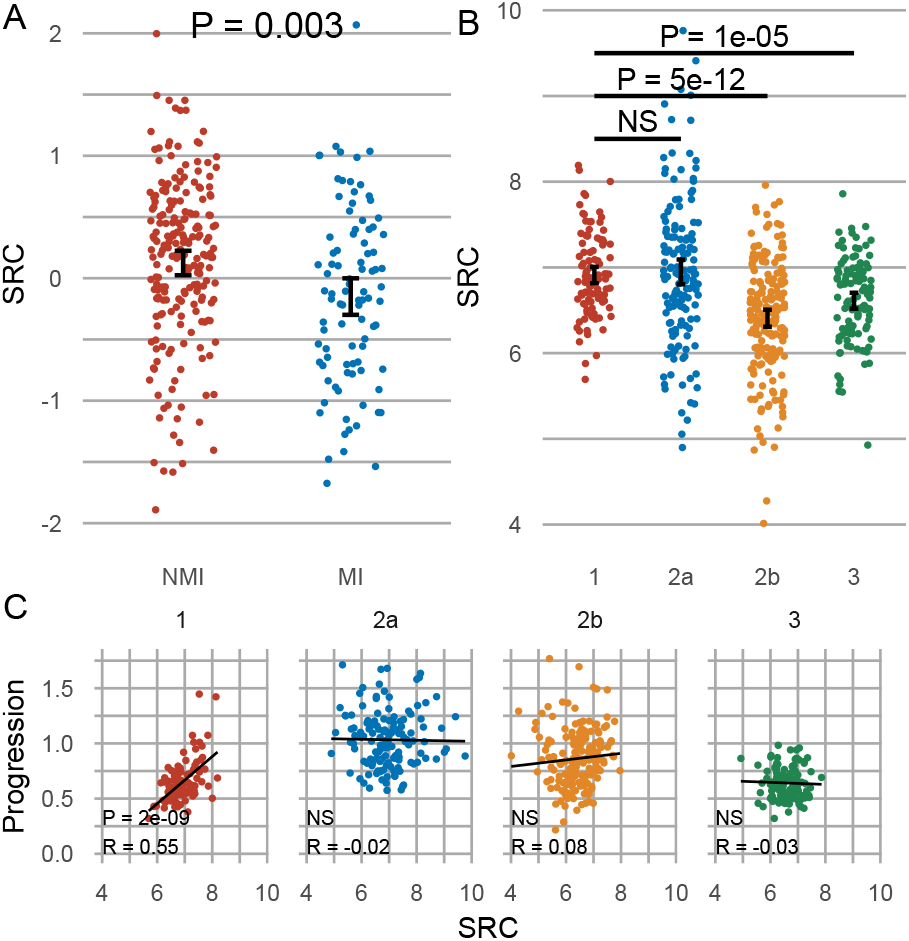
SRC is enriched in less aggressive bladder cancer subtypes. **A:** SRC expression in the Lund cohort, stratified by muscle invasion. **B:** SRC expression in the UROMOL 2021 cohort, stratified by UROMOL 2021 class. **C:** Pearson correlation between progression score and SRC expression. NMI: Non-muscle invasive, MI: Muscle invasive. CI = 95%.

Analyses of MIBCs revealed similar patterns between subtype aggressiveness and SRC expression, with SRC RNA (Figure 2, A) and protein expression (Figure 2, B) enriched in luminal papillary and luminal subtypes versus the others, and total and active site phosphorylated (Y416) SRC protein levels were also selectively enriched in the least aggressive luminal-papillary TCGA subtype.

**Figure 2:**
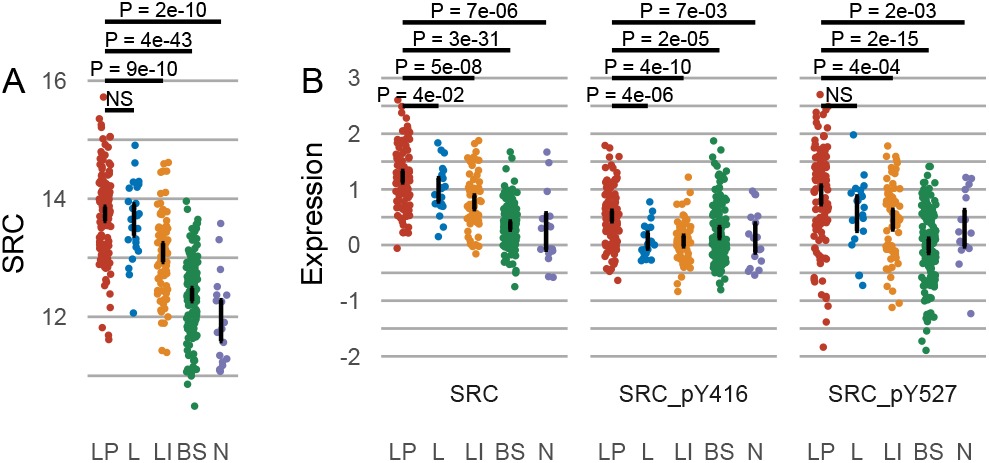
SRC is enriched in luminal muscle invasive bladder cancer. **A:** SRC expression in the TCGA cohort, stratified by TCGA subtype. **B:** Src protein and phosphorylated species expression in the TCGA cohort, stratified by subtype. LP: Luminal papillary, L: Luminal, LI: Luminal infiltrated, BS: Basal squamous, N: Neuronal. CI = 95%.

### Differential SRC expression in human cell lines

We next examined whether SRC expression was also heterogeneous in a panel of 30 human bladder cancer cell lines. To compare to human tumors, we first classified the cells by consensus class^14^. We found that our luminal-papillary (LP) cell lines expressed significantly higher levels of SRC than the basal squamous (BS) lines (Figure 3, A), consistent with our findings in human tumors. On the other hand, using GSVA to calculate gene set enrichment scores across classes, we found that the EMT signature was significantly upregulated in the BS lines when compared to the LP lines (Figure 3, B).

**Figure 3:**
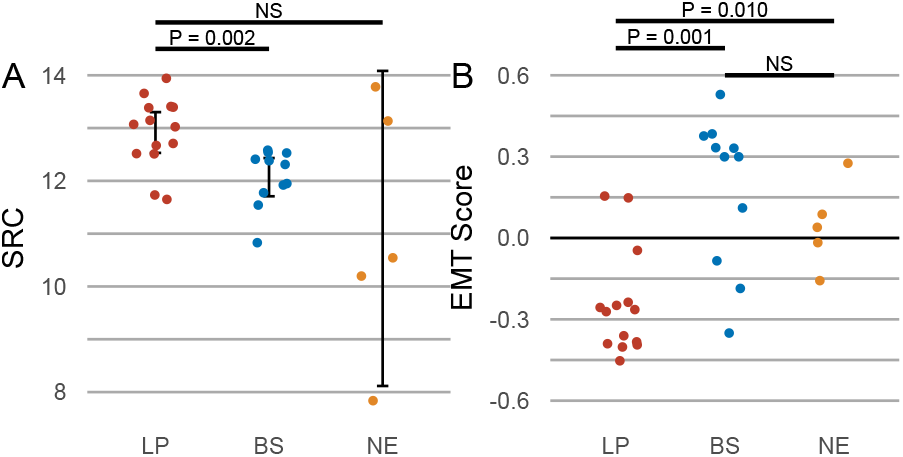
Bladder cancer cell lines recapitulate tumor subtypes. **A:** SRC expression in cell lines stratified by consensus classifier. **B:** Hallmark geneset enrichment scores stratified by consensus subtype. Shown Hallmarks are those with statistically significant ANOVA scores. LP: Luminal papillary; BS: Basal squamous; NE: Neuroendocrine-like. CI = 95%.

### Differential cell line motility across consensus class

To determine if motility differed across consensus classes, we performed two–dimensional migration assays using uncoated transwells, and invasion assays using Matrigel coated transwells using cells from within each class. We noted that LP lines tended to migrate (Figure 4, C) and invade (Figure 4, D) less readily than

**Figure 4:**
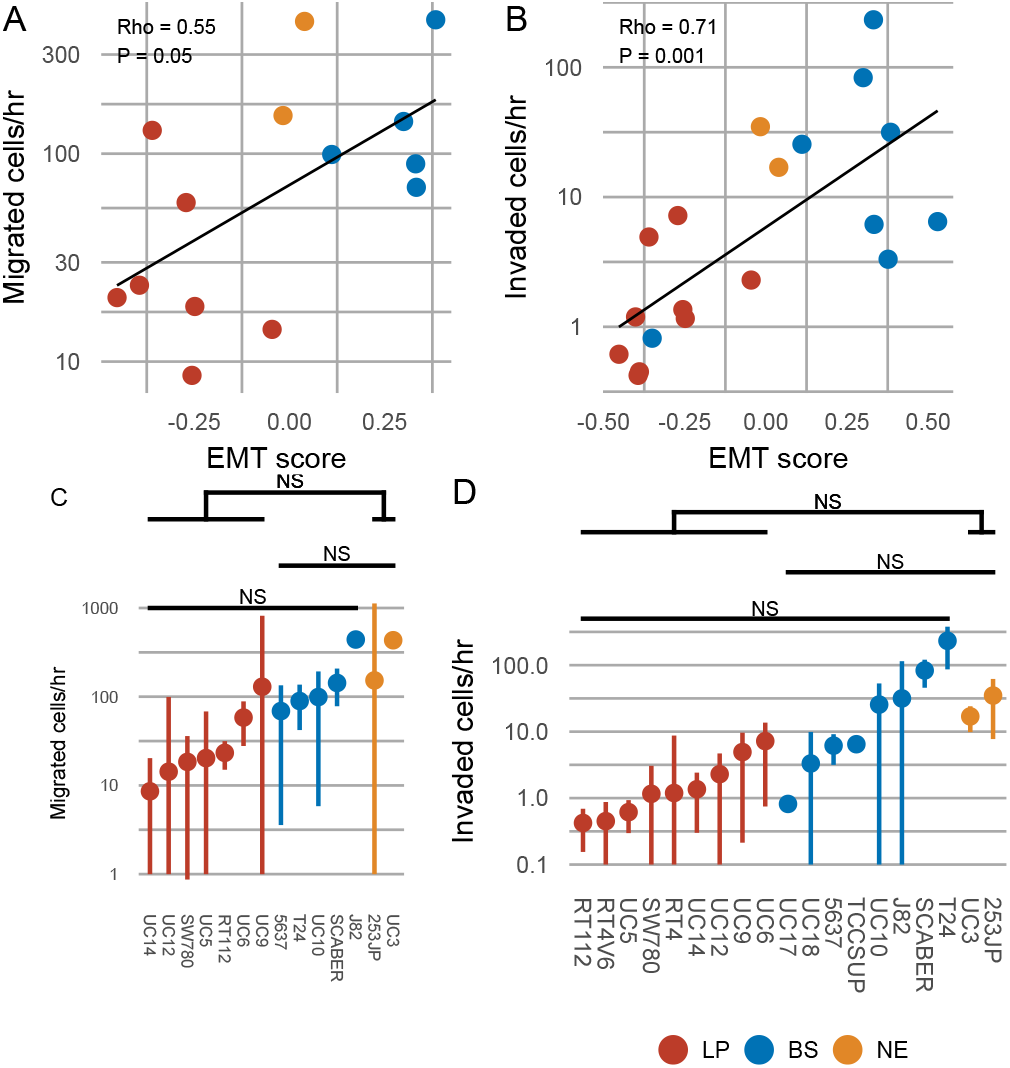
Motility across consensus subtypes. Migration (**A, C**) and invasion (**B, D**) rates of cell lines in Boyden chamber assays, correlated by EMT hallmark signature (**A, B**) or stratified by consensus subtype (**C, D**). LP: Luminal papillary; BS: Basal squamous; NE: Neuroendocrine-like. NS not significant. CI = 95%.

BS lines, but the differences were not statistically significant. However, we did note a statistically significant correlation between EMT score and both migration (Figure 4, A) and invasion (Figure 4, B).

### Differential sensitivity to SRC across consensus class

We next determined if sensitivity to Src inhibition varied across consensus classes using migration assays. We exposed representative LP and BS lines to the Src inhibitor bosutinib. On average, LP lines migrated at 53% their original rate, while BS lines migrated 72% their average rate, with individually statistically significant reductions in RT112 (p = 0.0028) and SCaBER (p = 0.00106) (Figure 5).

**Figure 5:**
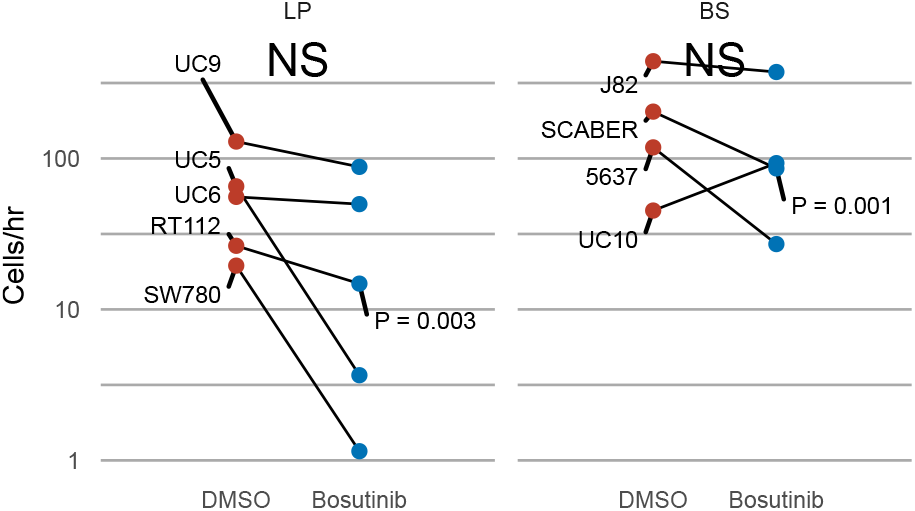
Sensitivity to Src inhibition across consensus subtypes. Changes in migration rates across consensus subtypes without (DMSO, -) or with (+) 1μM bositinib. LP: Luminal papillary; BS: Basal squamous

We additionally examined the effects of the chemically-distinct inhibitor, saracatinib (AZD0530). Although bosutinib had no effect on UM-UC6 migration, saracatinib strongly inhibited migration in both RT112 and UM-UC6 (Figure 8). Because saracatinib also inhibits the TGF-β receptor^15^, we tested the effects of galunisertib, a TGF-β receptor I inhibitor. Galunisertib had no effect on migration in RT112 but did appear to reduce rates consistently (but not significantly) in UC6 (Figure 8). The directionality of these effects implies migration may be driven more by TGF-β activity in UM-UC6, while migration may be driven more by Src in RT112.

We generated doxycycline-inducible SRC knockdown (SRC iKD) lines, validated by qPCR and Western blot (Figure 6, A) and performed migration assays with and without 48h 1μg/mL doxycycline pre-incubation. Similar to small molecule inhibitors, RT112 demonstrated an insignificant reduction in migration upon SRC KD, while UM-UC6 showed a modest increase in migration upon SRC KD (Figure 6). Because Src can also be required for proliferation^16^, we also measured the effects of SRC KD on proliferation upon in two cell lines. Bosutinib inhibited proliferation in both cell lines, though to a similar extent in all lines, implying that differences in migration seen with bosutinib are likely not driven by differences in proliferation. A more targeted approach with SRC knockdown showed a consistent but insignificant reduction in proliferation only in UM-UC6 (Figure 9).

**Figure 6:**
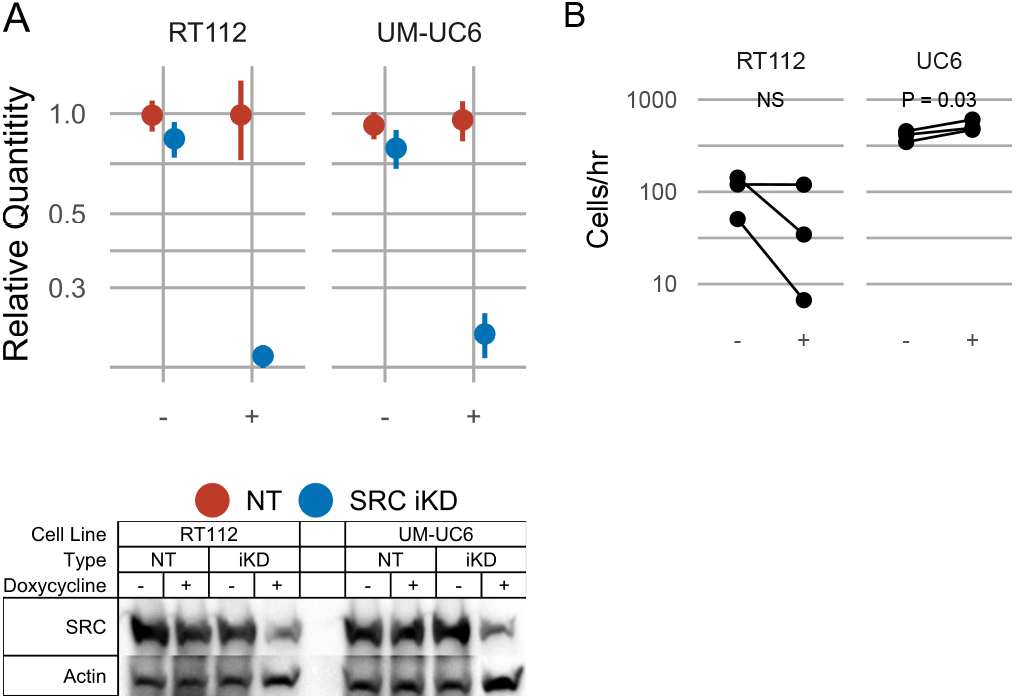
Src knockdown has differential effects on migration across cell lines. **A**: Src knockdown efficiency, measured by ΔΔCt qPCR (top), or Western blot (bottom) without (−) and with (+) 1μg/mL doxycycline, 48hr. **B** Migration rates of doxycycline-inducible SRC knockdown cell lines both without (−) and with (+) 48hr 1μg/mL doxycycline pre-incubation.

### *in vivo* effects of SRC inhibition

Finally, to directly examine the effects of SRC inhibition on metastasis, we pre-incubated the RT112 and UM-UC6 iKD cells +/-doxycycline before inoculating them into the tail veins of immunodeficient NSG mice housed with or without doxycycline in their drinking water. RT112 produced metastases in the lungs, liver, and spine, resulting in failure to void and hind leg paralysis, whereas UM-UC6 caused lymph node and lung metastases. SRC knockdown extended survival in mice inoculated with UM-UC6 but had no effect in animals with RT112 metas-tases (Figure 7) and did not affect the sites of metastasis in either model.

**Figure 7:**
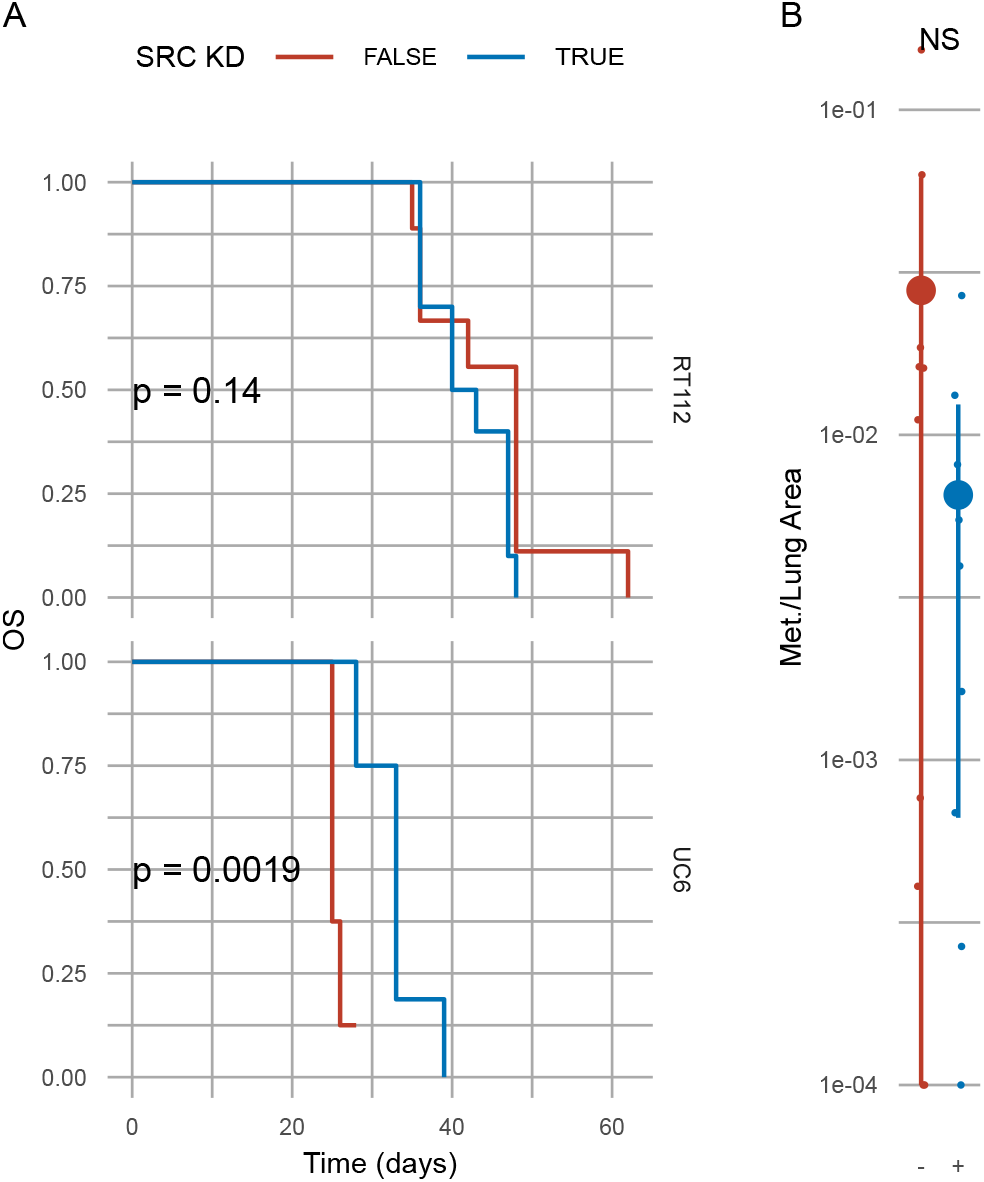
Src knockdown has differential *in vivo* effects across cell lines. **A:** Overall survival of NSG mice tail-vein injected with conditional knockdown cell-lines, with or without doxycycline drinking water. **B:** Ratio of UM-UC6 lung metastasis area to lung area, as quantified by ImageJ, in mice exposed to (+) or not exposed to (−) doxycycline. CI = 95%

**Figure 8:**
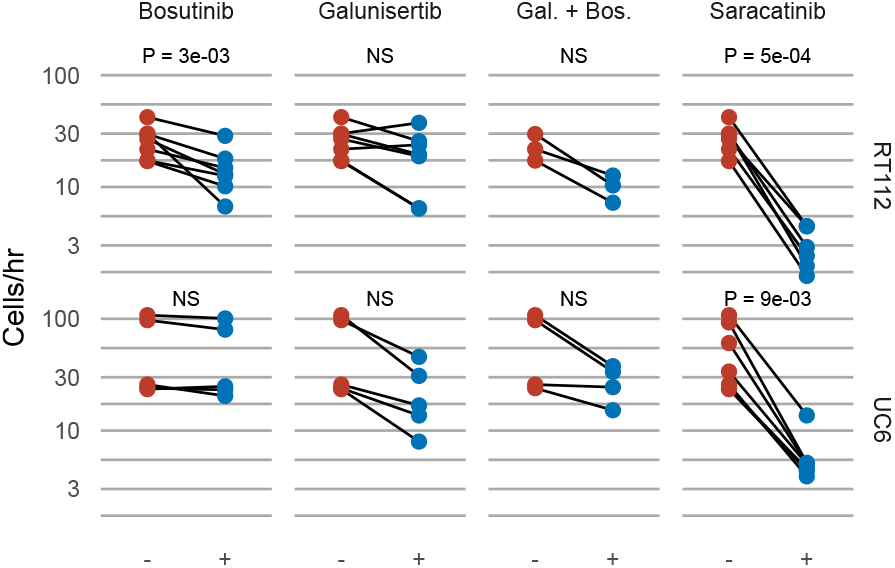
UM-UC6, not RT112, migration more sensitive to TGFβR I inhibition than Src inhibition. Migration rates of cells exposed to DMSO (−) or 1μM bosutinib, 1μM galunisertib, 1μM galunisertib + 1μM bosutinib, or 1μM saracatinib (+).

**Figure 9:**
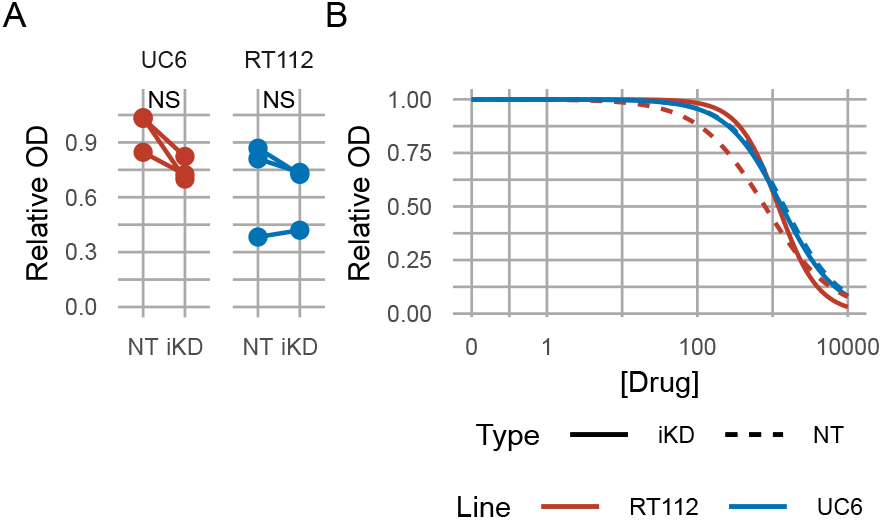
UM-UC6 proliferation may be inhibited by SRC knockdown. **A:** MTT assay of UM-UC6 and RT112 conditional SRC (iKD) or non-targeting (NT) knockdown cells exposed to 1μg/mL doxycycline. **B:** UC6 and RT112 exposed to increasing concentrations of bosutinib (nM).

## DISCUSSION

Src was one of the first discovered proto-oncogenes^2^, and over 100 years of study have revealed it serves a complex and varied role in cancer^16^. In this study, we sought to clarify the paradoxical role of Src as a metastasis suppressor in bladder cancer by testing the hypothesis that its effects may be dependent on molecular subtype. Overall, we observed modest and inconsistent effects of chemical SRC inhibitors or conditional SRC knockdown in conventional two-dimensional invasion and migration assays, and SRC knockdown also had modest effects in a model of experimental metastasis. SRC inhibition did appear to moderately inhibit proliferation in UM-UC6 cells, but not RT112, which may explain why knockdown extended survival in the UM-UC6 metastasis model. It is possible that established human cell lines and two-dimensional tissue culture are not adequate models of primary human tumors. On the other hand, our results reinforce the original conclusion, that SRC is enriched in luminal papillary human primary tumors. The luminal papillary subtype is associated with earlier state disease and better prognoses than are tumors belonging to the other molecular subtypes. In particular, luminal papillary tumors are characterized by gene expression signatures associated with terminal differentiation, suggesting that SRC may play an important role in this process. We also plan to directly test this possibility in appropriate preclinical models in future studies.

## SUPPLEMENTARY

## MATERIALS & METHODS

### Chemicals and reagents

Saracatinib (Selleck, S1006), bosutinib (Selleck, S1014), and galunisertib (MedChemExpress, HY-13226) were dissolved in DMSO at stock concentrations of 10mM and stored at −80°C. Prior to use, each inhibitor was diluted in culture medium yielding a maximum DMSO concentration of 0.1%.

### Cell culture

All parental cell lines were obtained from the Pathology Core of the Bladder Cancer SPORE at MD Anderson Cancer Center, with the exceptions of T24 and SCaBER where were purchased from the American Type Culture Collections. Cells were cultured as monolayers with Minimum Essential Medium (Gibco, 11095080) containing 10% fetal bovine serum (Corning, 35-011-CV), 1% nonessential amino acids (Gibco, 11140050), 1% vitamin solution (Gibco, 11120052), 1% penicillin-streptomycin solution (Gibco, 15140122), and 1% sodium pyruvate (Gibco, 11360070). Cells were incubated 37°C in a humidified, 5% CO2 atmosphere. Cell line identity confirmed by DNA finger-printing through the MD Anderson Characterized Cell Line Core and the Johns Hopkins School of Medicine Genetic Resources Core Facility. These experiments were performed with mycoplasma positive cells.

### Generation of conditional knockdown cell lines

Conditional knockdown lines were generated using SMARTvector Inducible Lentiviral shRNAs (Horizon Discovery). Lines were transduced either with lentivirus with constructs containing shRNA designed to target SRC (V3IHSPGG_10847004) or a non-targeting control (VSC6580) with a PGK RNA Pol II promoter. UM-UC6 was seeded at 10,000 cells/mL and RT112 at 30,000 cells/mL. Transduction was performed without serum, and transduced cells selected with 2.5μg/mL puromycin (InvivoGen, ant-pr-1).

### Transwell migration and invasion assays

Subconfluent cells were trypsinized, resuspended in serum containing medium, centrifuged, washed with PBS, centrifuged, and resupsended in serum free medium. 30,000 to 60,000 cells were then seeded into the upper chamber of the transwell in 0.5ml of serum free medium. The lower chamber was loaded with 30% FBS-containing medium to establish a chemo-attractant gradient. Migration or invasion was allowed to proceed for 4 to 20 hours. Membranes were then fixed, uninvaded cells wiped away, and remaining cells stained with gentian violet or DAPI. The entire filter was imaged by light microscopy and invaded cells quantified with CellProfiler^17^. 8µm pore Matrigel coated transwells (Corning, 354480) and uncoated transwells (Corning, 354578) were purchased from Corning.

### Western blotting

Subconfluent cells were harvested in RIPA buffer containing protease (cOmplete, Roche 11836153001) and phosphatase (PhosSTOP, Roche, 4906837001) inhibitors. Following protein quantification by BCA assay (Thermo Scientific, 23235), samples were diluted and boiled in 2x 2-mercaptoethanol containing Laemmli sample buffer (BioRad, 1610737). SDS-PAGE gel electrophoresis was run at 100 volts for 1.5 hours in Tris-Glycine-SDS buffer (BioRad, 1610732), followed by transfer onto nitrocellulose membrane in Tris-Glycine buffer (BioRad, 1610734) with 20% methanol for 2 hours at 100 volts. Membranes were blocked in casein blocking buffer (Sigma-Aldrich, B6429) for 1 hour, then incubated for 2 hours or overnight in primary antibody (1:1000 or 1:5000 dilution, 0.1X casein), washed in TBS-T (1X TBS, BioRad, 1706435; 0.1% Tween 20, Sigma-Aldrich, P7949), and incubated in secondary antibody for 1 hour (1:5000 dilution, 0.1X casein). In instances where multiple proteins of similar molecular weight were probed, identical but separate blot were performed at the same time. Loading control (actin) was examined on previously probed blots. Primary antibody Src was purchased from Cell Signaling (2123, RRID:AB_2106047, diluted 1:1000). Actin was purchased from Sigma-Aldrich (A2066, RRID:AB_476693, diluted 1:5000). Secondary antibodies were purchased from LI-COR (LI-COR 926-32210, 926-32211, diluted 1:5000).

### Tail Vein Injection

NSG mice were warmed in a mouse restrainer using a heat lamp to facilitate vasodilation. The tail was sterilized with 70% ethanol. A 100 µL cell suspension in PBS, containing 2×10^6 UM-UC6 cells or 5×10^5 RT112 cells, was slowly injected into the tail vein after confirming blood backflow.

### Lung Metastasis Quantification

All physical H&E slides were scanned using the CONCENTRIQ platform for metastasis quantification. The tumor fraction in the lungs was quantified by calculating the ratio of the total metastatic area to the total lung area using FIJI^18^.

### Statement on Animal Care and Welfare

The Johns Hopkins University Institutional Animal Care and Use Committee approved the experimental procedures used in this study (approval no. MO16M463) on December 2021. All animal housing and experiments were conducted in strict accordance with the institutional Guidelines for Care and Use of Laboratory Animals at Johns Hopkins University.

## Statistical Analyses

All analyses were performed using R^19^ (v. 4.4.1). P < 0.05 was used as a cutoff for statistical significance. T-tests are two-tailed. Paired t-tests were used to compare drug/control within biological replicates. Log-rank test was used to test for survival differences (survival^20^). GSVA^21^ was used to calculate signature enrichment scores.

## Data Availability

All data and code used in this paper are publicly available. Code used to obtain data and generate figures can be found at https://github.com/McConkeyLab/2024_aragaki-src. The Lund cohort data were obtained from GEO (GSE32894). The UROMOL cohort data were obtained from the supplementary and source data from the original paper^13^. TCGA BLCA data were obtained from the Genomic Data Commons, using the pipeline provided at https://github.com/McConkeyLab/general_tcga. Cell line RNA expression was obtained from our publically available package, cellebrate: https://github.com/McConkeyLab/cellebrate.

## References

1. Rous, P. A SARCOMA OF THE FOWL TRANSMISSIBLE BY AN AGENT SEPARABLE FROM THE TUMOR CELLS. Journal of Experimental Medicine 13, 397–411 (1911)

2. Martin, G. S. The road to Src. Oncogene 23, 7910– 7917 (2004)

3. Yeatman, T. J. A renaissance for SRC. Nature Reviews Cancer 4, 470–480 (2004)

4. Thomas, S. et al. Src and Caveolin-1 Reciprocally Regulate Metastasis via a Common Downstream Signaling Pathway in Bladder Cancer. Cancer Research 71, 832–841 (2011)

5. Wu, Y. et al. Src phosphorylation of RhoGDI2 regulates its metastasis suppressor function. Proceedings of the National Academy of Sciences 106, 5807–5812 (2009)

6. Kamat, A. M. et al. Bladder cancer. The Lancet 388, 2796–2810 (2016)

7. Robertson, A. G. et al. Comprehensive Molecular Characterization of Muscle-Invasive Bladder Cancer. Cell 171, 540–556 (2017)

8. Damrauer, J. S. et al. Intrinsic subtypes of highgrade bladder cancer reflect the hallmarks of breast cancer biology. Proceedings of the National Academy of Sciences 111, 3110–3115 (2014)

9. Choi, W. et al. Identification of Distinct Basal and Luminal Subtypes of Muscle-Invasive Bladder Cancer with Different Sensitivities to Frontline Chemotherapy. Cancer Cell 25, 152–165 (2014)

10. Knowles, M. A. & Hurst, C. D. Molecular biology of bladder cancer: new insights into pathogenesis and clinical diversity. Nature Reviews Cancer 15, 25–41 (2014)

11. Cappellen, D. et al. Frequent activating mutations of FGFR3 in human bladder and cervix carcinomas. Nature Genetics 23, 18–20 (1999)

12. Fanning, P. et al. Elevated expression of pp60csrc in low grade human bladder carcinoma. Cancer research 52, 1457–1462 (1992)

13. Lindskrog, S. V. et al. An integrated multi-omics analysis identifies prognostic molecular subtypes of non-muscle-invasive bladder cancer. Nature Communications 12, (2021)

14. Kamoun, A. et al. A Consensus Molecular Classification of Muscle-invasive Bladder Cancer. European Urology 77, 420–433 (2020)

15. Klaeger, S. et al. The target landscape of clinical kinase drugs. Science 358, (2017)

16. Thomas, S. M. & Brugge, J. S. CELLULAR FUNCTIONS REGULATED BY SRC FAMILY KINASES. Annual Review of Cell and Developmental Biology 13, 513–609 (1997)

17. Stirling, D. R. et al. CellProfiler 4: improvements in speed, utility and usability. BMC Bioinformatics 22, (2021)

18. Schindelin, J. et al. Fiji: an open-source platform for biological-image analysis. Nature Methods 9, 676– 682 (2012)

19. R Core Team. R: A Language and Environment for Statistical Computing. (2024)

20. Therneau, T. M. A Package for Survival Analysis in R. (2024)

21. Hänzelmann, S., Castelo, R. & Guinney, J. GSVA: gene set variation analysis for microarray and RNA-Seq data. BMC Bioinformatics 14, 7 (2013)

